# Single-Cell Multiomics Defines Tolerogenic Extrathymic Aire-Expressing Populations with Unique Homology to Thymic Epithelium

**DOI:** 10.1101/2021.11.05.467513

**Authors:** Jiaxi Wang, Caleb A. Lareau, Jhoanne Bautista, Alexander Gupta, Katalin Sandor, Joe Germino, Yajie Yin, Matt Arvedson, Gabriella C. Reeder, Nathan T. Cramer, Fang Xie, Vasilis Ntranos, Ansuman T. Satpathy, Mark S. Anderson, James M. Gardner

## Abstract

The Autoimmune Regulator (*Aire*) gene, well defined for its role in medullary thymic epithelial cells (mTECs) and immune self-tolerance, is also expressed in extrathymic *Aire*-expressing cells (eTACs) in the secondary lymphoid organs. eTACs have been shown to be hematopoietic antigen presenting cells (APCs) and potent inducers of immune tolerance (1–3). However, the precise identity and function of these cells remain unclear. Here, we use high-dimensional single-cell multiomics and functional approaches to define eTACs at the transcriptional, genomic, and proteomic level. We find that eTACs consist of two similar cell types: CCR7+ Aire-expressing migratory dendritic cells (AmDCs) and a unique Aire-hi population co-expressing Aire and RAR-related orphan receptor gamma-t (RORγt). The latter, which have significant transcriptional and genomic homology to migratory dendritic cells (migDCs) and mTECs, we term Janus cells (JCs). All eTACs, and JCs in particular, have a highly accessible chromatin structure and high levels of broad gene expression, including tissue-specific antigens, as well as remarkable transcriptional and genomic homology to thymic medullary epithelium. As in the thymus, Aire expression in eTACs is also dependent on RANK-RANK-ligand interactions. Furthermore, lineage-tracing shows that JCs are not precursors to the majority of AmDCs. Finally, self-antigen expression by eTACs is sufficient to mediate negative selection of T cells escaping thymic selection and can prevent autoimmune diabetes in non-obese diabetic mice. This transcriptional, genomic, and functional symmetry between a hematopoietic Aire-expressing population in the periphery and an epithelial Aire-expressing population in the thymus suggests that a core biological program may influence self-tolerance and self-representation across the spectrum of immune development.

## INTRODUCTION

The Autoimmune Regulator (*Aire*) gene plays an essential role in the maintenance of central tolerance, and its function has been extensively defined in thymic medullary epithelial cells, where, among other roles, it regulates the expression of a diverse range of otherwise tissue-specific self-antigens (4, 5). Aire expression has also been observed in the secondary lymphoid organs in both mice and humans (6, 7), and self-antigen expression in such extrathymic Aire-expressing cells (eTACs) is sufficient to cause deletion or inactivation of cognate T cells (1, 2). Recently, eTACs have also been demonstrated to be essential for the maintenance of normal maternal-fetal immune tolerance (8). Thus, the precise identity and biology of these extrathymic Aire-expressing populations is of substantial biological and clinical interest.

While Aire expression has been described in various dendritic cell populations (1,2,7,9– 12), the precise identity of extrathymic Aire-expressing cells has remained elusive. Recent reports have suggested that the principal Aire-expressing population in the secondary lymphoid organs may be a subset of type-3 innate lymphoid cells (ILC3s), based largely on the co-expression of Aire and RAR-related Orphan Receptor, gamma T (RORγt) (3), encoded by the Rorc gene and known for its essential role in T helper cell type 17 (Th17) and ILC3 differentiation. A growing body of evidence has also shown that RORγt-expressing populations characterized as ILC3s can play important roles as APCs to promote immune tolerance in a range of contexts, most notably in commensal tolerance in the gut (13, 14). Furthermore, single-cell transcriptional mapping of murine DC lineages has also recently identified populations of RORγt expressing DCs (15). Thus, the identification of an Aire- and RORγt-coexpressing migratory DC-like population may have implications for some of these divergent findings.

Recent evidence suggests that migratory dendritic cells may have important regulatory functions in the maintenance of immune tolerance (16–18). Initially characterized by their participation in the generation of memory immune responses to acquired antigens (19, 20), migratory DCs have more recently been suggested to participate in the regulation and repression of immune responses. Transcriptional characterization of migDCs by the ImmGen consortium suggested enrichment for an immunomodulatory and tolerogenic transcriptional program (21). Deletion of migDC populations, or interference with their migration, worsened autoimmunity in a range of mouse models (16, 17). More recently, populations of tumor-associated migratory DCs (termed mregDCs) were described which had both tolerogenic phenotype and function, and regulated immune responses to neoplasia (18). However, the signals and transcriptional regulatory circuits driving such tolerogenic phenotypes in migDCs remain obscure.

Here, we use high-throughput single-cell multiomics and a range of functional approaches to define eTACs as migratory DC-like populations, and identify among them a new mixed-phenotype Aire and RORγt-expressing population which we term Janus cells (JCs). We show that these populations have uniquely high degrees of broad chromatin accessibility and gene expression, including a range of tissue-specific antigens and immunomodulatory transcripts. Self-antigen expression in eTACs also leads to self-tolerance among T cells that escape thymic selection. Further, we find that eTACs have remarkable transcriptional and genomic homology to medullary thymic epithelium, despite being hematopoietic in origin. This convergent transcriptional circuitry between thymic and peripheral Aire-expressing populations, despite their distinct stromal and hematopoietic origins, may provide unique insights into the fundamental mechanisms maintaining self-tolerance throughout immune development.

## RESULTS

### Extrathymic Aire-expressing cells consist of distinct populations of migratory DC-like cells

To define the identity and basic biology of extrathymic Aire-expressing populations in an unbiased fashion, we sought to characterize these cells utilizing high-dimensional single-cell multiomics, including single-cell RNA sequencing (scRNA-seq) and ATAC with Selected Antigen Profiling by sequencing (ASAP-seq). We enriched for rare populations in the lymph nodes (LNs) by magnetic-column depletion of T and B cells from the digested, pooled LNs of WT and Aire-reporter Adig mice (1). Next, enriched cells were either directly subjected to parallel multiome sequencing (scRNAseq and ASAP-seq) from WT mice, or flow-sorted for all live, GFP+ cells from Aire-driven IGRP-GFP (Adig) mice, a previously characterized Aire-reporter mouse strain (1) (**Fig. 1A, S1A**). From the scRNA-seq data (**Fig. S1B-C**), the un-sorted WT sample (n=6,973 cells) allowed us to generate a map of rare and common secondary lymphoid populations, while the GFP-sorted sample (n=2,532 cells) from Adig mice allowed us to deeply profile eTAC populations and further map them into the immune landscape to define their identity. Using our combined data resource, we identified discrete populations of eTACs (GFP+ cells from the Adig LNs) which aligned closely with GFP and Aire transcript expression (**Fig. 1B, D**), confirming both sort purity as well as the fidelity of the GFP reporter. In the merged data, we were able to identify a broad range of rare lymphoid, myeloid, and stromal cell types, and found eTACs represented two distinct populations, both of which shared high transcriptional similarity to migratory DCs (**Fig. 1C-G, S2A-B**). The majority of Aire expression was seen in migratory DC1/2s, and we referred to these populations as Aire-expressing migratory dendritic cells (AmDCs), although these AmDCs did not cluster separately as a distinct group from Aire-negative migDCs (**Fig. 1C**). In contrast, the eTAC subset with the highest Aire expression was a discrete cluster, clearly distinct in RNA space after dimensional reduction, that expressed both high levels of *Aire* as well as RAR Related Orphan Receptor C (*Rorc*; **Fig. 1C, D, G**), a gene otherwise associated with innate and adaptive lymphocyte populations such as Th17 and ILC3s (22). Importantly, these same eTAC populations were also all present in reference (non-Aire-enriched) samples alone, though at lower frequency (**Fig. 1B, S1D**). Despite its Rorc expression, this AireHi population otherwise appeared more closely related to the migratory DC lineage in tSNE space, and expressed comparable levels of *Zbtb46*, *Ccr7*, and MHC class II. Because of the distinct phenotype and dual expression of AireHi/Rorc+, we refer to the cell population as Janus cells (JCs), in reference to the two-faced classical deity.

**Figure 1.**
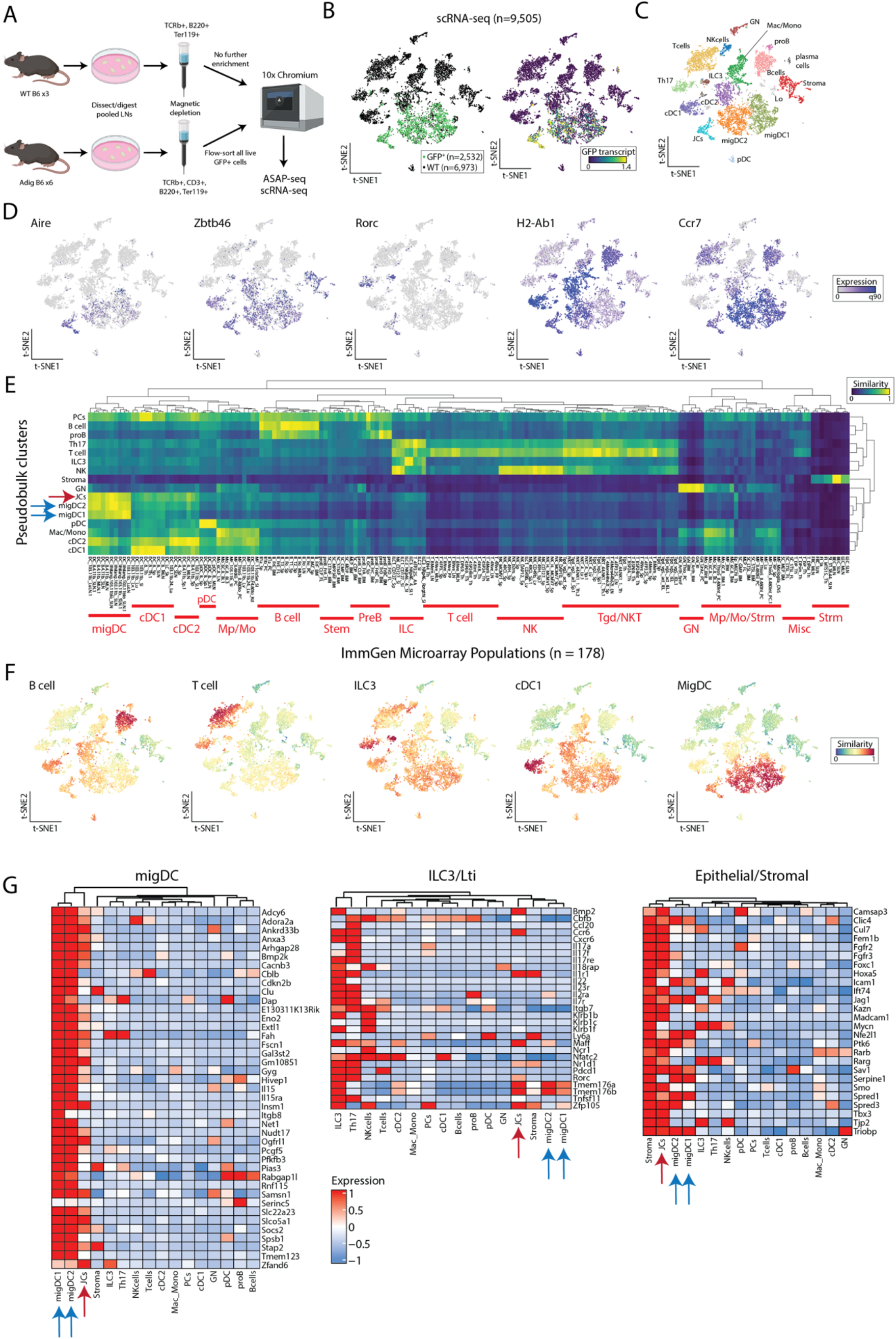
Extrathymic Aire-expressing cells consist of distinct populations of migratory DC-like cells. (A) Schematic of cell isolation and enrichment for scRNAseq and ASAPseq. (B) Reduced dimensionality representation of scRNA-seq data, indicating the GFP+ population and GFP transcript. (C) Annotated cell clusters of populations in scRNA-seq data. (D) Gene expression of selected transcripts used to define populations, including *Aire* and *Rorc*. (E) Hierarchical clustering of pseudobulk scRNA-seq clusters by all ImmGen microarray data using a scaled cosine similarity metric. Rows min-max normalized. Immgen populations cluster identities annotated in red text as indicated. Aire-expressing populations are indicated with blue (migDC1/2) and red (JC) arrows. (F) Single-cell cosine similarity scores for scRNA-seq data using indicated ImmGen populations. (G) Gene expression heatmaps for selected gene sets, z-score-normalized per row. Aire-expressing populations are indicated with blue (migDC1/2) and red (JC) arrows.

To more precisely assign identity to extrathymic Aire-expressing populations in an unbiased fashion based on their entire transcriptome rather than individual marker genes, we next performed unsupervised two-dimensional hierarchical clustering and cosine similarity scoring of each pseudo-bulked population against the entire Immunologic Genome Project (ImmGen) microarray database of 178 annotated reference immune populations (23) in RNA space (**Fig. 1E**). Our informatics approach verified clustering of eTAC populations (JCs with AmDCs), as well as high transcriptional similarity to reference ImmGen populations collectively identified as migratory DCs (23). Reciprocally, we then utilized select ImmGen reference populations to assign cosine similarity scores on a per-cell basis to each cell profiled by scRNA-seq, which again demonstrated that both JCs and AmDCs had high transcriptional similarity to migratory DCs (**Fig. 1F, S2B**). These unbiased, reciprocal approaches to cell identity assignment indicate that all eTAC populations appear to be most transcriptionally similar to migratory dendritic cells.

Noting that migDCs have been characterized by a common canonical transcriptional signature (21), we verified this signature was highly expressed in all Aire-expressing clusters (JCs and migDC1/2) (**Fig. 1G**). And despite recent claims (3), we found that JCs clustered broadly with myeloid and not lymphoid lineages. JCs share less homology with defined populations of either NKp46+ or LTi ILC3s and do not express detectable levels of most canonical ILC3-family transcripts (*Il17a/f, Il22, Il23r*) (24). However, some genes, including *Ccr6* and *Tmem176a/b*, were shared between the populations. Interestingly, eTACs in general, and JCs in particular, also transcriptionally resembled cells in the epithelial/stromal lineage (**Fig. 1G**), suggesting some transcriptional similarity to non-hematopoietic populations.

### Single-cell-chromatin accessibility supports the genomic identity of eTACs as myeloid populations with uniquely accessible chromatin

To further characterize eTACs at both the genomic and proteomic level, we subjected the same populations of cells purified from Aire-reporter and WT mice described above (**Fig. 1A**) to combined single-cell ATAC and surface protein multiomics (ASAP-seq) (25). In strong concordance with the single-cell RNA sequencing data, we again found that *Aire* accessibility was largely confined to migDCs and JCs (**Fig. 2A-C**). To independently validate the cellular identity of these populations based on their chromatin profiles, we next performed unsupervised two-dimensional hierarchical clustering and cosine similarity scoring of each pseudo-bulk population against the entire ImmGen ATAC-seq database that consisted of 89 populations (26). While the ImmGen ATAC database does not include migratory DC populations, JCs and migDCs again both co-clustered with each other and showed the highest cosine similarity to dendritic cells (**Fig. 2D,** red and blue arrows). Conversely, these eTAC populations showed low similarity to ILCs or other lymphoid lineages and were also similar to medullary thymic epithelial cells (**Fig. 2D,** green arrow). These identities were reciprocally validated using per-cell cosine similarity projections based on accessible chromatin (**Fig. S3A**).

**Figure 2.**
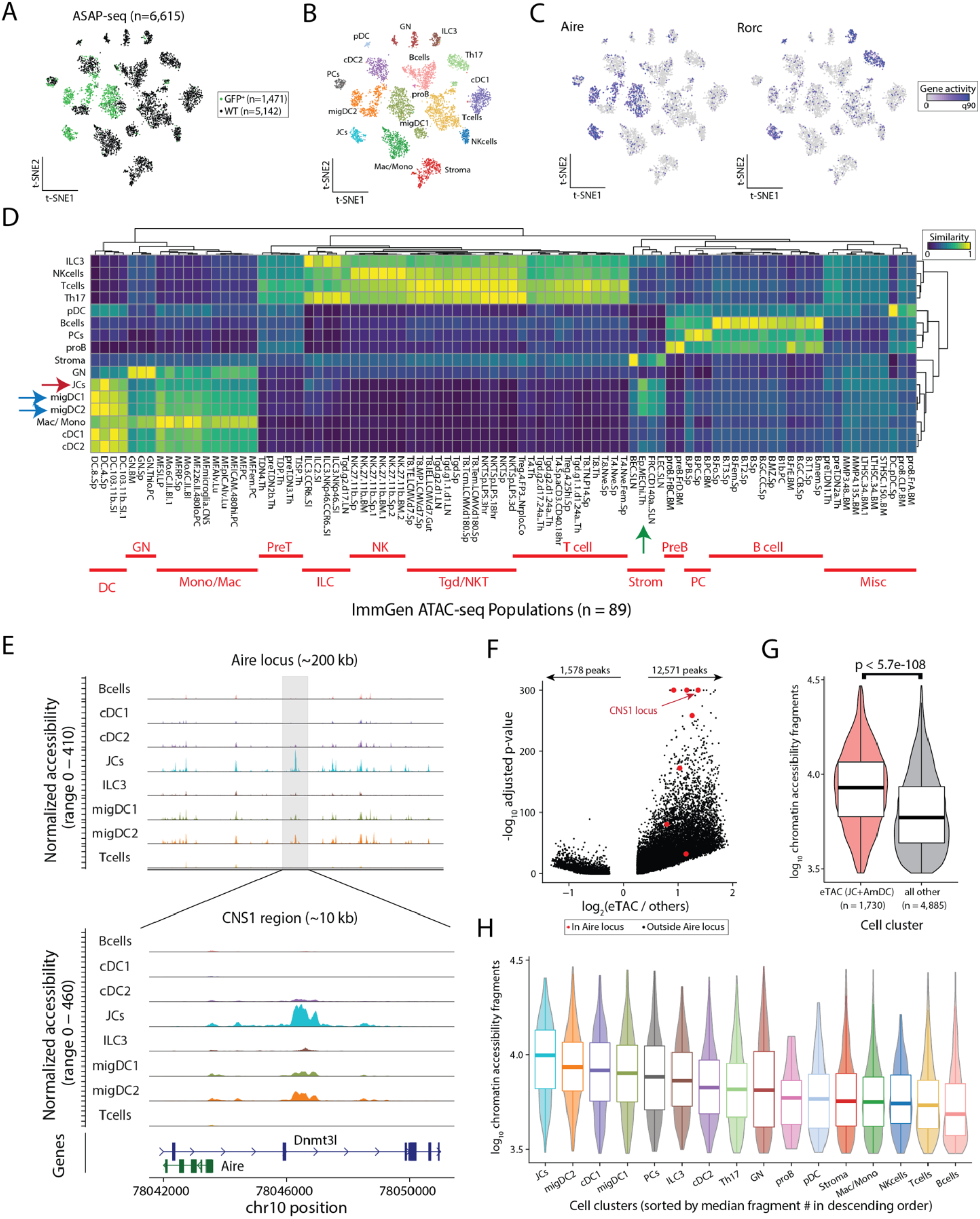
Single-cell-chromatin accessibility supports the genomic identity of eTACs as myeloid populations with uniquely accessible chromatin. (A) Reduced dimensionality representation of ASAP-seq data, indicating the GFP and WT sample origins. (B) Annotated cell clusters of populations in ASAP-seq data. (C) Gene activity scores of *Aire* and *Rorc* across ASAP tSNE. (D) Hierarchical clustering of pseudobulk ASAP-seq clusters by all ImmGen ATAC-seq data using a scaled cosine similarity metric. Rows min-max normalized. Immgen populations cluster identities annotated in red as indicated. Aire-expressing populations are indicated with blue (migDC1/2) and red (JC) arrows, and mTECs from the ImmGen database with a green arrow. (E) Accessible chromatin landscape near the *Aire* locus, including the previously described CNS1 region (bottom). (F) Volcano plot of differential accessibility peaks, indicating the number of peaks with greater (n=12,571) or less (n=1,578) accessibility in Aire expressing populations. *Aire* locus peaks in large red dots; CNS1 locus as indicated. (G) Comparison of number of accessible chromatin fragments between eTACs and non-eTAC populations in the LN by box/violin overlay. Statistical test: Mann-Whitney test. (H) Per-cluster abundance of accessible chromatin fragments/cluster, box/violin overlay, noting highest accessibility in JCs. Boxplots: center line, median; box limits, first and third quartiles; whiskers, 1.5× interquartile range.

Mapping the accessibility of the cis-regulatory region defining the Aire locus in these populations showed distinct peaks in JCs and migDCs in the CNS1 region, known to be essential for *Aire* expression (27, 28) (**Fig. 2E**). Notably, our analysis of this locus also revealed several additional peaks unique to Aire-expressing populations that had not been previously described (**Fig. 2E, Fig. S3B**). Interestingly, global differential accessibility analyses of eTACs against all other Aire-negative populations showed significantly increased chromatin accessibility in Aire-expressing cells relative to all other LN populations, including at the *Aire* locus (**Fig. 2F, G**). To support that this was not due to batch effects, we also evaluated the same chromatin accessibility between eTACs and other LN populations in the WT dataset alone and again observed significantly increased accessibility among eTACs (**Fig. S3C**). To confirm this result was not due to skewing from a particular low-accessibility population, we also examined the per-cell chromatin accessibility of each individual population and found the same trend—JCs had the highest chromatin accessibility of any cell type in the LN, followed closely by migDC1/2s (**Fig. 2H**). Overall, our analyses support a marked increase of chromatin accessibility in eTACs, a striking observation given Aire’s known role in chromatin regulation in the thymus (29–31) and in governing the expression of a broad range of tissue-specific antigens.

### Focused multiomic analysis of eTAC populations identifies broad transcriptional upregulation and antigen presentation in all eTACs with uniquely high TSA expression in JCs

To better define the biology and relationships of eTAC subsets, we next focused specifically on GFP-sorted populations. We again confirmed that eTACs consist principally of AmDC1/2s and JCs, as well as small numbers of cells from B-cell and ILC lineages (**Fig. 3A, Fig. S4A**). Coincident with their increased chromatin accessibility, eTACs also had high levels of global gene expression compared with all other populations (**Fig. S4B**), with JCs having the highest level of global gene expression among eTAC clusters (**Fig 3B, Fig. S4B)**. Differentially upregulated genes in JCs included a wide range of transcripts including both *Aire* and *Rorc*, as well as a distinct set of unique JC-specific genes (**Fig. 3B, Table S1**). Among these JC-specific genes, we identified a large number of transcripts previously classified as tissue-specific antigens (TSAs) (32) (**Fig. 3C, Table S1**). Indeed, when we surveyed the global TSA expression, we found that JCs expressed the highest numbers of TSA genes, both when compared against AmDCs and more globally against all LN populations, presumably because of their overall increased levels of gene expression (**Fig. 3D, Fig. S4B, C**) and greater chromatin accessibility. Interestingly, these TSAs were particularly enriched for neuronal-associated and germ cell/placental antigens (**Fig. S4D**), suggesting both potential disease models in which to interrogate the function of these populations, as well as potential links to the established findings that loss of eTACs leads to a breakdown in maternal-fetal tolerance (8).

**Figure 3.**
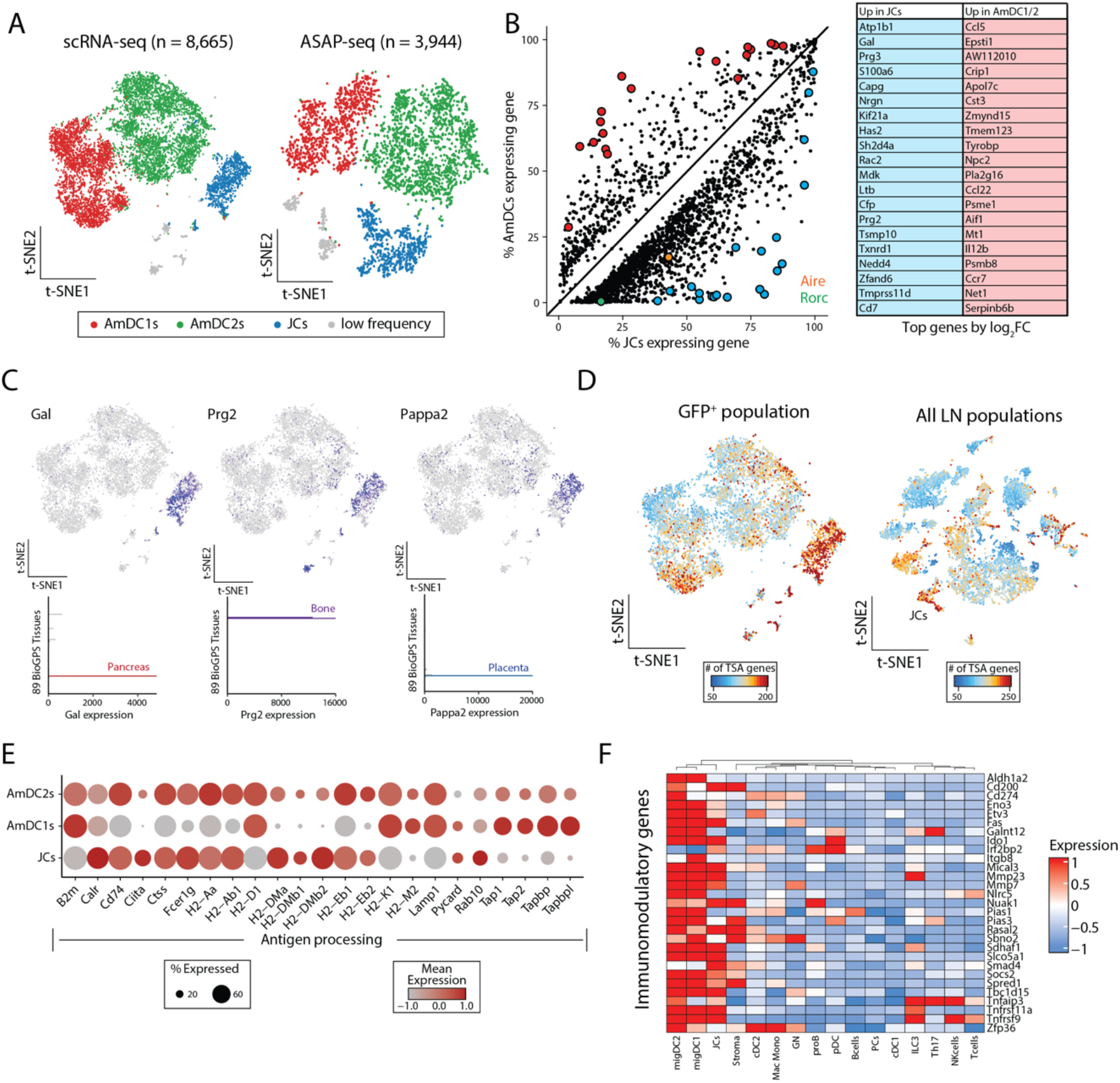
Focused multiomic analysis of eTAC populations identifies broad transcriptional upregulation and antigen presentation in all eTACs with uniquely high TSA expression in JCs. (A) Reduced dimensionality representation for all GFP+ scRNA-seq and ASAP-seq data. (B) Differential gene expression analyses of JCs (x-axis) and AmDCs (y-axis), with *Aire* (orange) and *Rorc* (green) highlighted. Top 20 genes are shown in the table on the right. (C) Expression of three tissue specific antigen genes, *Gal, Prg2,* and *Pappa2*, differentially upregulated in JCs. The top represents expression in the GFP+ scRNA-seq data and the bottom bar graphs represent tissue restricted expression as indicated from GeneAtlas tissue populations. (D) Number of tissue-specific antigen (TSA) genes for the GFP+ only (left) and all lymph node (right) populations, noting the relative abundance in JCs. (E) Dot plot of antigen processing and presentation gene set showing expression in GFP+ populations. (F) Heatmap of immunomodulatory genes across all lymph node populations from scRNA-seq data, z-score normalized per row.

Both JCs and AmDCs also expressed a diverse array of genes associated with antigen processing and presentation in both the class I and class II MHC pathways (**Fig. 3E**). Additionally, eTACs also expressed a range of genes associated with immune tolerance and tolerogenic dendritic cell maturation (21, 33), including *Socs2, Fas, Cd200, Ido1, Smad4,* and *Irf2bp2* (**Fig. 3F**). These findings suggest that these populations may be involved in both the expression and/or acquisition of self-antigens, and supports the existing evidence (1–3) that eTACs may promote antigen-specific immune tolerance among interacting lymphocyte populations.

### Surface-marker multiomics combined with functional flow cytometry and lineage-tracing allows for identification and characterization of eTACs and their lineage relationships

Although JC and AmDC subpopulations of eTACs appear transcriptionally similar, the lineage relationship between them was unclear. As single-cell ASAP-seq allows for identification of a broad range of surface-markers associated with each population, we next sought to define the surface proteins unique to AmDCs and JCs, which would allow us to distinguish and trace these populations. Differential analysis defined several surface proteins relatively more prevalent in either AmDCs (CD11c, CD45, MHCII) or JCs (CD200; **Fig. 4A, B**).

**Figure 4.**
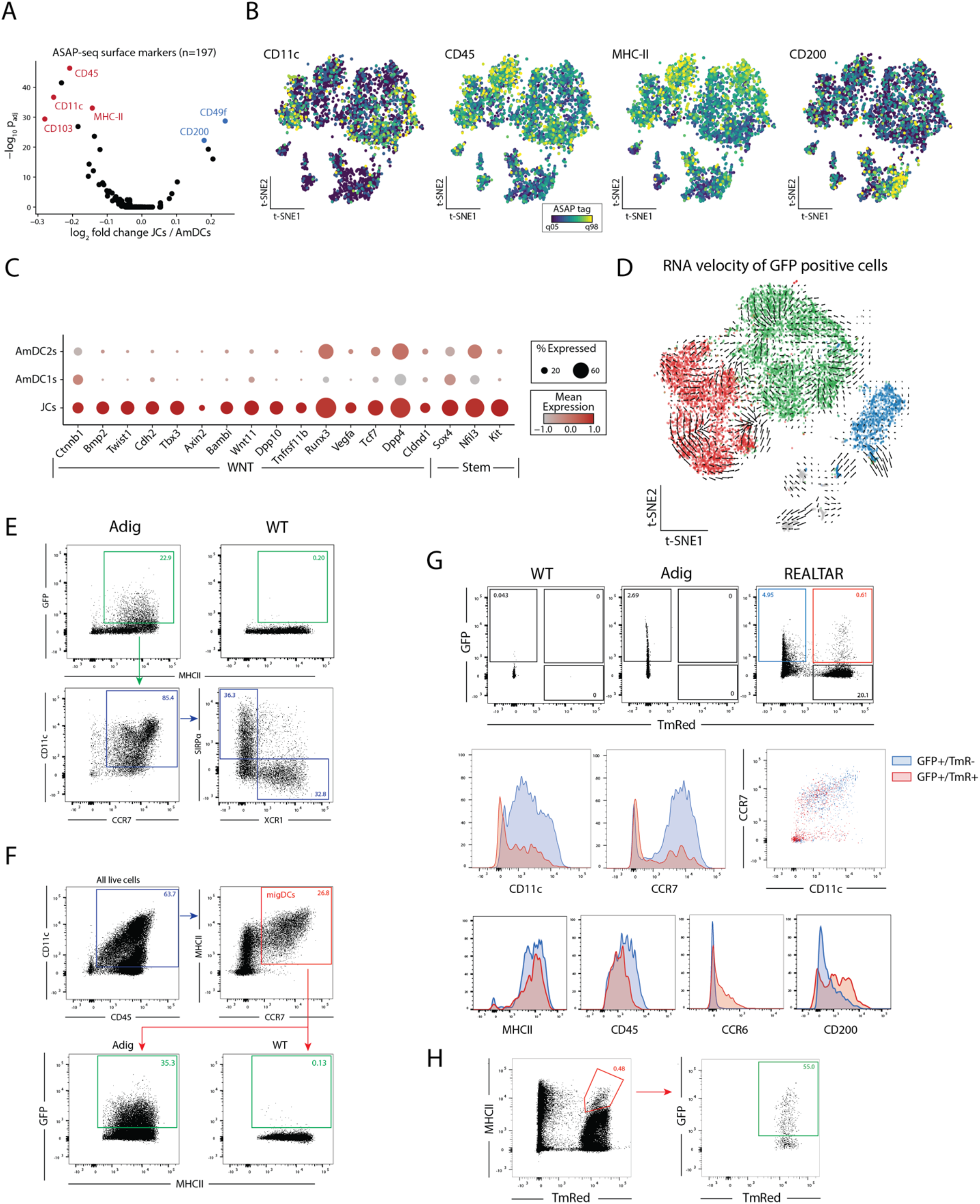
Surface-marker multiomics combined with functional flow cytometry and lineage-tracing allows for identification and characterization of eTACs and their lineage relationships. (A) Volcano plot of surface markers measured by ASAP-seq comparing JCs to AmDCs. (B) Per-cell visualization of selected surface markers indicated in panel A. (C) Dot plot summarizing expression of WNT and Stem-like genes over-expressed in JCs compared to Aire-expressing migratory DCs. (D) RNA velocity analysis of GFP+ populations. (E) Flow cytometry from WT and Adig lymph nodes, pre-gated on single, live, and dump (TCRbeta, CD19, SiglecF, F4/80, NK1.1, Ly6C)-negative cells. (F) Flow cytometry from WT and Adig lymph nodes, pre-gated on single and live cells. (G) Flow cytometry from WT, Adig, and REALTAR mice lymph nodes, pre-gated on single, live, and dump (TCRbeta, CD19, SiglecF, F4/80, NK1.1, Ly6C)- negative cells. (H) Flow cytometry from REALTAR mouse lymph nodes, pre-gated on single and live cells.

Furthermore, transcriptional data from the scRNAseq analysis suggested that JCs were highly enriched for expression of a range of Wnt-signaling pathway and progenitor-associated factors previously correlated with stem-like cell fates (34–38) (**Fig. 4C**). Interestingly, Wnt signaling through e-cadherin and beta-catenin also has a role in maintaining tolerogenic function in dendritic and myeloid cells (39, 40). To determine whether JCs’ progenitor-like phenotype reflected their status as a precursor to AmDCs, we utilized both bioinformatic and functional approaches. Notably, RNAvelocity (41) analysis showed no evidence of developmental vectors projecting from JCs to AmDCs (**Fig. 4D**). To verify this inference, we sought to characterize these relationships between respective eTAC populations by genetic lineage tracing and flow cytometry. First, using the Adig mouse, we verified that the majority of eTACs consist of migratory DC1/2s in roughly equal proportions of the two populations (**Fig. 4E**) – consistent with proportions found in our multiome data. Surprisingly, assessed as a percentage of total migDCs, AmDCs represent over 30% of the total migDC pool at baseline (**Fig. 4F**).

Next, to better identify JCs by flow cytometry and understand their lineage relationship to AmDCs, we used the previously described RORγt-Cre mouse (42). RORγt-Cre mice were crossed with Adig Aire-reporter and Rosa26-LSL-TmR mice to generate RORγt-Expression And Lineage-Tracer and Aire Reporter (REALTAR) mice. The REALTAR mouse allowed us to identify cells actively transcribing Aire by GFP expression, and cells expressing or having ever expressed RORγt by TmRed expression. Using this model, we could clearly identify two populations: a larger, distinct population of GFP+/TmR- cells, as well as a smaller population of GFP+/TmR+ cells (**Fig. 4G**). Surface-marker characteristics of these populations matched what we observed by scRNA-seq and ASAP-seq (**Fig. 3B, 4A-B**), that JCs (GFP+/TmR+) expressed dramatically lower levels of CD11c and CCR7 than AmDCs (GFP+/TmR-; **Fig. 4G**). This was validated by staining for a number of other differential markers between AmDCs and JCs including MHCII, CD45, CCR6, and CD200 (**Fig. 1G, 4A, 4G**), which are consistent with previously published results (3).

Thus, we confirmed that JCs have a unique CCR7lo/CD11clo surface-marker phenotype distinct from AmDCs, despite expression of both genes at the transcript level in JCs. This lineage-tracer system also validates that JCs express the RORγt isoform of Rorc. Further, these data suggest that the majority of AmDCs are not lineage-traced by RORγt and thus do not appear to arise from an RORγt-expressing JC precursor. Finally, it is interesting to note that among MHCII-hi RORγt lineage-traced (TmRed+) cells, over 50% of this population is in fact Aire-expressing (GFP+) (**Fig. 4H**). This suggests that perhaps JCs should be considered in the interpretation of previously described tolerogenic phenotypes attributed to MHCII-expressing ILC3s using RORγt-Cre systems (13, 14).

### eTACs are defined by transcriptional and regulatory homology to thymic medullary epithelium

While the global transcriptional and functional profile of eTACs suggested a high similarity with migratory DCs, we noted that our cosine-similarity scoring against reference ImmGen populations demonstrated significant transcriptional homology to another population – medullary epithelial cells of the thymus (mTECs; **Fig. 5A, B).** Interestingly, this was particularly pronounced among JCs. Indeed, the top differentially expressed genes defining JCs were highly enriched only in mTECs when assessed among all other annotated populations in the ImmGen RNA-seq database (**Fig. 5C**). To define the expression of these genes more precisely relative to Aire expression in the thymus, we aggregated an atlas of prior published scRNA-seq data on thymic epithelial cells and performed integrative analyses using scVI (43) to establish a dimensionally-reduced map of thymic epithelial populations across the full spectrum of their development. The integrated data was annotated according to the population identities defined in the source data (44–47) (**Fig. S5**). In addition to being expressed in mTECs, many of these JC-specific genes appeared tightly associated with Aire expression during mTEC development (**Fig. 5D**). Importantly, the majority of these genes were not, themselves, Aire-regulated in the thymus (three of the top 50 DE genes, 5.5% overall; **Table S2**) (48), but rather may represent Aire-associated transcriptional pathways with relevance for the biology and function of Aire in these populations. Strikingly, differential transcription factor binding-site accessibility analysis of the scATAC data demonstrated that the chromatin landscape of JCs was most highly enriched for transcription factor binding sites known for their roles driving Aire expression in mTECs, including RelB and NFkB, as well as less well-characterized HIVEP genes (**Fig. 5E**). While some of this overlap might be explained by binding-site sequence degeneracy, a number of these gene transcripts including the noncanonical NFkB-pathway genes *Relb* and *Nfkb2*, and *Hivep1* and *3*, appeared to be highly expressed in JCs and/or migDCs (**Fig. 5F**). Surveying the ImmGen ATAC database showed that these same transcription factor binding sites are also highly enriched in the open chromatin of medullary thymic epithelium (**Fig. 5G**, red arrow). Thus, our analyses indicate that JCs share a canonical transcriptional and accessible chromatin program with mTECs, likely mediated by Aire expression.

**Figure 5.**
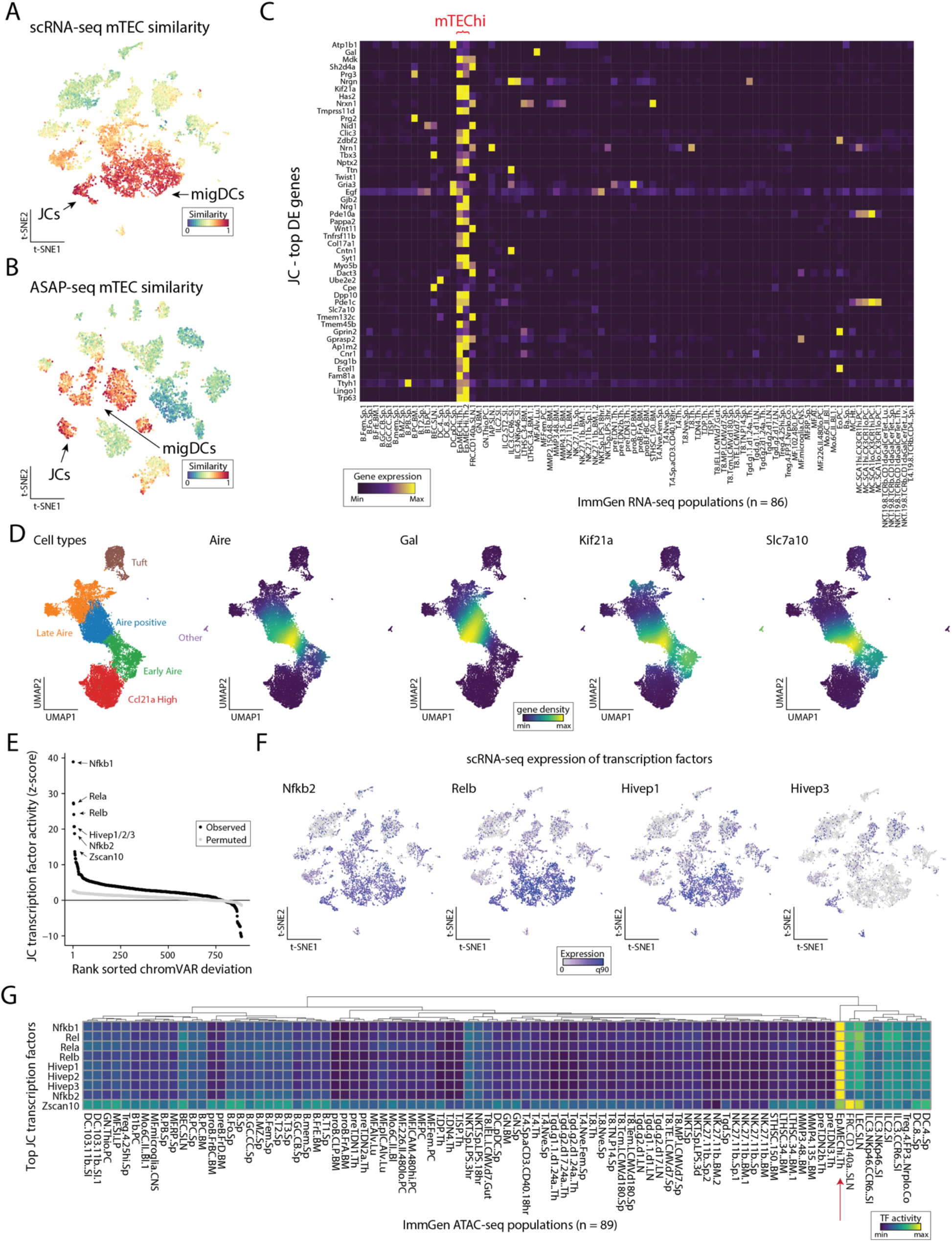
eTACs are defined by transcriptional and genomic homology to thymic medullary epithelium. (A) ImmGen similarity scores for scRNA-seq and (B) ASAP-seq data for mTEC populations. JCs and migratory DCs clusters for each embedding are noted by arrow. (C) Heatmap of ImmGen bulk RNA-seq data for genes most overexpressed in JCs relative to all other lymph node populations. mTEChi populations indicated in red text on top. (D) UMAP of aggregated published scRNA data from thymic epithelium showing annotated TEC subsets (left) and gene-density expression for *Aire* and genes enriched in JCs (right). (E) Rank-ordering of transcription factor motifs enriched in JCs compared to other populations. (F) Expression of transcription factors from panel E in scRNA-seq data. (G) ImmGen bulk ATAC-seq chromVAR deviation scores for all 89 populations, highlighting the mTEC population with red arrow. Rows represent top transcription factors identified from JCs and are min-max normalized.

### RANK-RANK(L) signaling is required for Aire expression in eTACs

Given the strong genomic and transcriptional evidence for RANK/NFkB signaling in eTACs and the well-established role of RANK-RANKL signaling in driving Aire expression in the thymus (49, 50), we next asked whether these signals are also required for Aire expression in eTACs. First, we found that RANK (Tnfrsf11a) expression was highly upregulated in eTACs. OPG (Tnfrsf11b), a soluble RANK antagonist that often serves in a negative-feedback regulatory mechanism in cells receiving high levels of RANK signaling (51, 52), was quite specifically expressed in JCs among peripheral immune populations (**Fig. 6A**). As previously demonstrated (46,53,54), inhibition of RANK signaling by antibody-blockade of RANKL led to a significant loss of Aire expression among mTECs in the thymus (**Fig. 6B, D**). The same was true in the periphery, where eTACs (all GFP+ cells) decreased in relative and absolute abundance after RANKL-blockade (**Fig. 6C, D**). Most notably, JCs (GFP+ CCR7-CD11c-) almost entirely disappeared (or lost Aire expression), suggesting that as in the thymus, Aire-expression in JCs depends on RANK/NFkB signals. This result again demonstrates the symmetry between Aire expression in the thymus and periphery, provides valuable insight into signals required to induce Aire expression in hematopoietic populations, and suggests that studies utilizing RANK blockade, for example in the setting of tumor immunity (54, 55), should also consider the impact on extrathymic populations.

**Figure 6.**
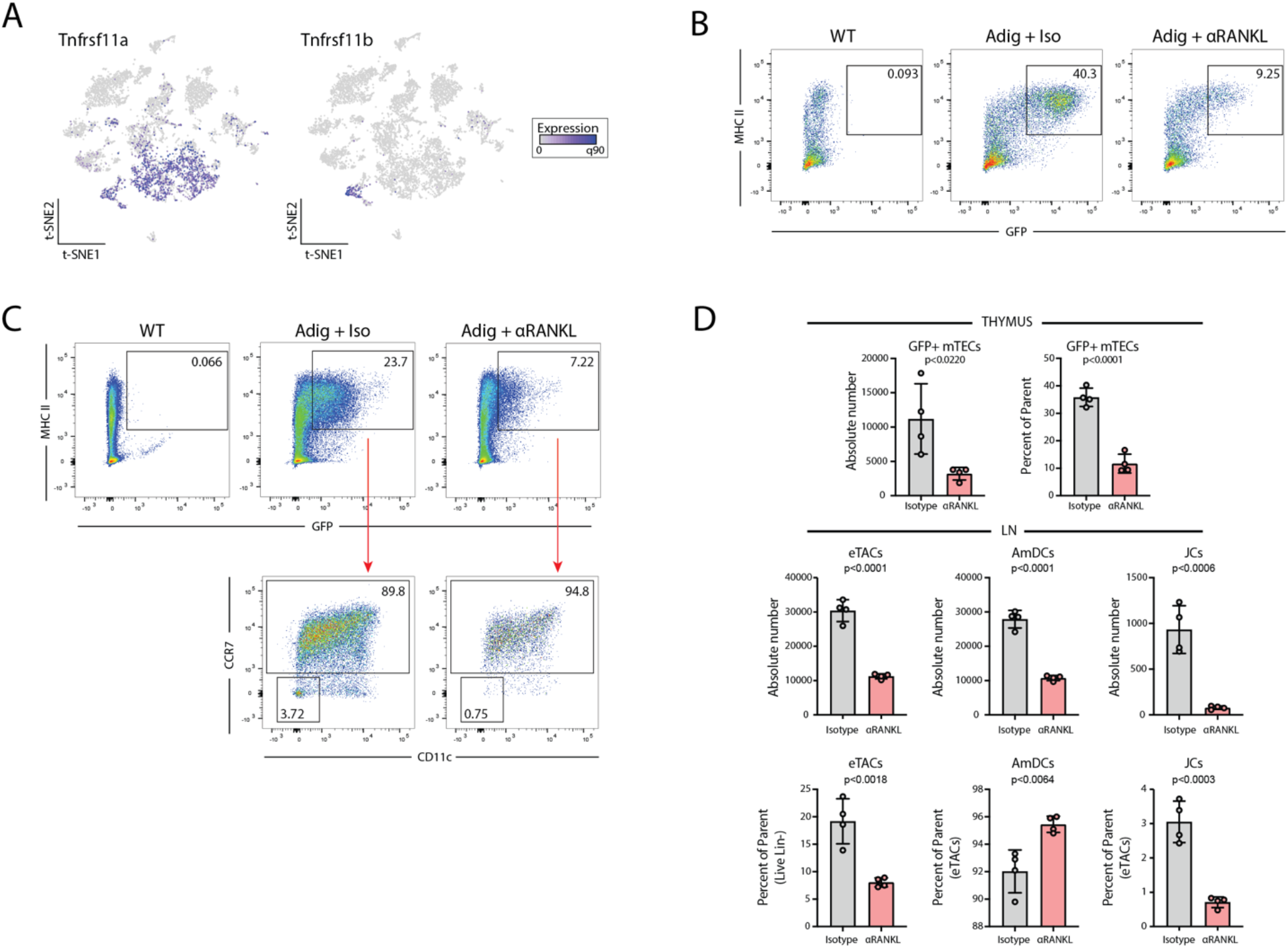
RANK-RANK(L) signaling is required for Aire expression in eTACs. (A) Gene expression of RANK (*Tnfrsf11a*) and OPG (*Tnfrsf11b*) from scRNA-seq data. (B) Flow cytometry of Percoll-enriched thymic epithelial cells from WT and Adig mice treated with isotype or αRANKL, pre-gated on single, live, CD11c-, CD45-, EpCAM+ cells. (C) Flow cytometry of lymph nodes from WT and Adig mice treated with isotype or αRANKL treatment, pre-gated on single, live, and dump (TCRbeta, CD19, SiglecF, F4/80, NK1.1, Ly6C)-negative cells. (D) Quantitation of percentage and absolute cell numbers from B and C, unpaired t-test.

### Pancreatic self-antigen expression in eTACs is sufficient to induce deletion of T cells escaping thymic selection, and to prevent autoimmune diabetes

Given these transcriptional and genomic similarities between mTECs and eTACs, despite their different origins, we asked whether self-antigen presentation by eTACs is sufficient to mediate tolerance for T cells that escape thymic negative selection. To assess this, we utilized a feature of the Adig mouse, which also expresses a pancreatic self-antigen, islet-specific glucose-6-phosphatase related protein (IGRP), under the control of the Aire promoter (1). In order to generate mice in which only eTACs and not mTECs expressed IGRP, we thymectomized WT and Adig non-obese diabetic (NOD) mice and simultaneously transplanted wildtype thymi under the kidney capsule, then sub-lethally irradiated and reconstituted them with bone marrow from the 8.3 mouse, a CD8+ TCR-transgenic specific for IGRP (56) (**Fig. 7A**). Despite concern that peripheral antigen might traffic back to the transplanted thymus and mediate thymic negative selection, we saw no evidence of this, as FACS analysis and tetramer staining of lymphocytes from these mice demonstrated no thymic negative selection and no loss of tetramer avidity (**Fig. 7B-D**). By contrast, in the secondary lymphoid organs, we observed profound deletion of IGRP-specific CD8+ T cells and significantly decreased tetramer avidity of surviving CD8+ T cells (**Fig. 7B-D**). Importantly, despite the large numbers of IGRP-specific T cells leaving the thymus in this permissive TCR-transgenic system, eTAC-mediated deletion in the secondary lymphoid organs was sufficient to entirely prevent these mice from developing diabetes (**Fig. 7E**).

**Figure 7.**
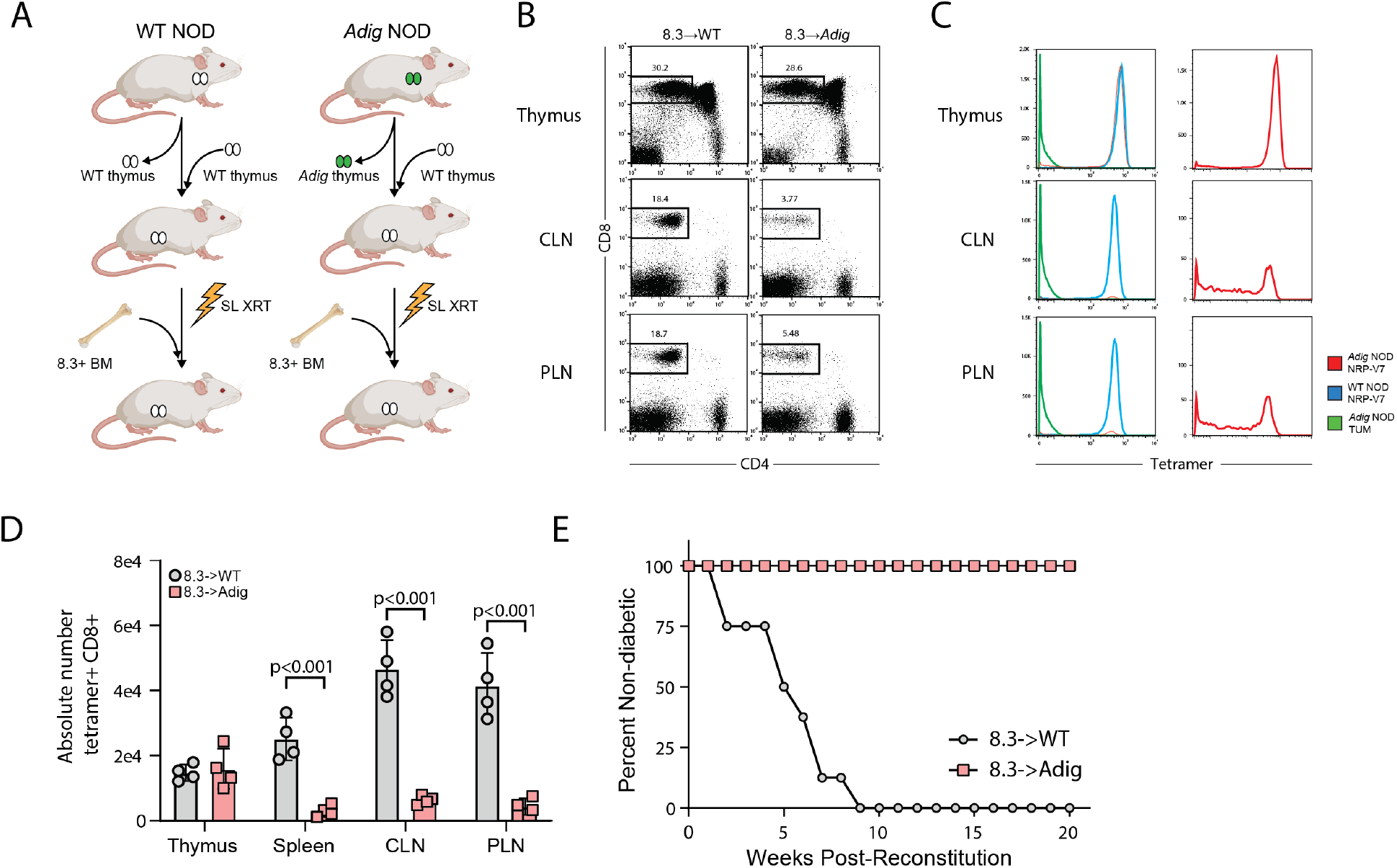
Pancreatic self-antigen expression in eTACs is sufficient to induce deletion of T cells escaping thymic selection, and to prevent autoimmune diabetes. (A) Experimental design for thymic swap and BM chimerism in NOD and Adig NOD mice. SL XRT: sublethal irradiation. BM: bone marrow. (B) Flow cytometry showing lymphocyte populations from indicated tissues. CLN: cervical lymph node. PLN: pancreatic lymph node. (C) Flow cytometry tetramer staining of lymphocyte populations from indicated tissues; panels on right show Adig NOD NRP-V7 staining alone to show distribution of avidity in the periphery, given small number of surviving cells. (D) Quantitation of absolute cell numbers from B, unpaired t-test. (E) Kaplan-Meier curves, diabetes-free survival after bone-marrow reconstitution with 8.3+ BM.

## DISCUSSION

A growing body of evidence has demonstrated that migratory DCs in the steady state may play a significant role in immune tolerance, with diverse examples in immune homeostasis, autoimmunity, and tumor immunity (16–18). Further, while traditional paradigms suggested an inverse correlation between DC maturity and immunogenicitiy, some mature DCs, particularly migratory DCs, acquire a discrete tolerogenic profile (33). However, the transcriptional signals driving such tolerogenic maturation has remained unclear. In parallel, work defining the identity and biology of eTACs has suggested a potential migDC-like phenotype (2, 9), but their precise identity and biology has remained unclear.

Here, we defined eTACs using high-dimensional multiomics as consisting of two distinct migratory DC-like populations, AmDCs and JCs, and show that self-antigen expression in eTACs is sufficient to cause deletion of autoreactive T cells escaping thymic selection and prevent autoimmune diabetes. Remarkably, we find a distinct transcriptional homology between eTACs and mTECs, and comparative analysis of the two populations identified a core transcriptional signature shared between the two populations. This suggests several promising future directions to define both the signals driving Aire expression and more broadly the transcriptional circuitry underlying self-education in the adaptive immune system. We show that eTACs share a common genomic accessibility signature with mTECs enriched in RANK/NFkB signaling pathways, and as in the thymus, RANK/RANKL signaling is required for Aire expression in eTACs. Beyond these known signals, several other pathways appear common to both populations, including Wnt-signaling and the HIVEP genes. These parallel analyses also suggest novel candidate regulatory regions around the Aire locus which may provide further insight into the regulation of Aire expression in both the thymus and periphery. Finally, the unique genomic accessibility and high transcriptional activity of eTACs generally, and JCs specifically, raises numerous fascinating questions about their biology and homeostasis, and the role of TSA expression within these populations.

While prior work had suggested that Aire/RORγt+ cells might be an ILC3-like population (3), here, we used unbiased whole-transcriptome and whole-chromatin single-cell profiling methods to strongly support the myeloid and DC-like phenotype of JCs. Their similarities to ILCs appear to be largely a product of RORγt-dependent gene expression, like *Tmem176a/b* (24). These AireHi/RORγt+ cells have previously been demonstrated to be capable of antigen processing and presentation (3), and we show these pathways are upregulated in both JCs and AmDCs. Combined with their highly accessible chromatin and their broad transcriptional program including a range of tissue-specific antigens, this suggests the population may play a role in immune self-education. Indeed, the representation of placental antigens among the TSAs expressed by JCs suggests a potential link to the recently reported requirement for eTACs in the maintenance of maternal-fetal tolerance in both syngeneic and allogeneic pregnancy (8). Interestingly, due to their lack of surface protein expression of either CD11c or CCR7, this population may have been overlooked in prior analyses of dendritic cell subsets. Further, a number of previous studies have described tolerogenic phenotypes and attributed peripheral regulatory T-cell (Treg) induction to MHCII-expressing ILC3s using RORγt-Cre/MHCII-flox systems (13, 14). The identification of RORγt-expressing JCs suggests that these cells should be considered as contributors to these phenotypes and deserve further study.

Numerous questions remain, including the contributions of peripheral Aire to normal immune homeostasis, the precise function of Aire within eTACs, and the mechanisms of eTAC-mediated tolerance induction. Additionally, while we have demonstrated here that JCs are most likely not precursors to the majority of AmDCs, JCs do share many phenotypic characteristics with migratory DCs and thus represent an intriguing discrete myeloid population whose developmental origin will be essential to further define. Similarly, the signals driving Aire expression in these populations, and in the thymus through shared programs, will require greater investigation to understand the physiologic significance of these regulatory networks. Finally, while eTACs have recently been demonstrated to be essential for normal immune homeostasis in maternal-fetal tolerance during pregnancy (8), the precise mechanisms of that tolerance remain unclear. While the expression of a range of pregnancy-associated and placental antigens in JCs offers a tantalizing clue, it remains to be demonstrated whether the observed phenotype is a direct result of this antigen reservoir loss, of antigen acquisition by these migratory populations, or of some other mechanism.

The remarkable parallelism between thymic and peripheral Aire-expressing cells does, however, suggest a uniquely conserved central and peripheral core circuitry for the maintenance of immune tolerance despite the divergent origin of these two cell types. Defining the biology of this core Aire-associated program may shed significant insight on basic Aire biology and its role in immune tolerance. Overall, work delineating the conserved transcriptional circuitry between these two tolerogenic populations could have broad relevance both for our understanding of normal immune homeostasis mechanisms, and for diverse applications in autoimmunity, maternal-fetal tolerance, tumor immunity, and transplantation.

## MATERIALS AND METHODS

### Mice

C57BL/6J (B6, H-2b), NOD, B6 RORγt-Cre, and B6 Rosa26-LSL-TmR mice were purchased from The Jackson Laboratory. B6 Aire-Driven IGRP-GFP (Adig) and NOD-Adig mice were generated by the lab and characterized as previously described (1). NOD Thy 1.1 8.3 TCR-transgenic mice, previously derived and characterized (56), were bred and utilized as bone marrow donors. All mice were housed in specific pathogen-free facilities at the University of California San Francisco, and all animal studies were approved by the Institutional Committee on Animal Use and Care at the University of California San Francisco.

### Antibodies and Tetramers

Fluorochrome- or biotin-tagged antibodies against the following mouse targets were purchased from: BD Biosciences – CD11c (N418), Siglec F (E50-2440); Thermo Fisher - TCRb (H57-597), CD3e (145-2C11), B220 (RA3-6B2), Ter119 (TER-119), MHCII (AF6-120.1); and BioLegend - CD4 (GK1.5), CD8 (53-6.7), NK1.1 (PK136), Ly6c (HK1.4), CD90.2 (30-H12), CD19 (6D5), F4/80 (BM8), CCR7 (4B12), XCR1 (ZET), SIRPa (P84), TCRgd (GL3), CD45 (30-F11), CD200 (OX-90), CCR6 (29-2L17), EpCAM (G8.8). Isotype control antibodies were purchased from same source as target-specific antibodies. Fixable Blue Dead Cell Stain Kit for UV excitation (Thermo Fisher) was used for viability staining for flow cytometry. DAPI (4′,6- diamidino-2-phenylindole) was used for viability staining for cell sorting.

MHC class I tetramers (I-Ag7) were provided by the NIH Tetramer Core Facility (Emory University, Atlanta, GA). I-Ag7 tetramers used for staining CD8+ T lymphocytes were the IGRP mimetope/class I MHC tetramer NRP-V7-PE (KYNKANVFL) or irrelevant mock tetramer TUM-PE (KYQAVTTTL).

### Thymectomy, thymic transplantation, bone marrow chimera generation

Wildtype neonatal (2do) B6 pups were used as thymic donors; thymi were procured and cultured in twelve-well transwell plates in 1.35mM 2-deoxyguanosine in complete DMEM-10 at 37 degrees for seven days. One day prior to transplant (on d6), 2-DG media was replaced with regular complete DMEM. At day seven of thymic incubation, 6-8-week-old NOD and Adig NOD mice were anesthetized with Ketamine/Xylazine and immobilized, and a midline submandibular incision made, the salivary glands were reflected upward, and a partial sternotomy was made. The thymic lobes were suction-aspirated, confirming two lobes aspirated per mouse, and the salivary glands reflected down to cover the partial sternotomy, and the skin closed with staples. The mice were then rotated on their sides, and the left flank shaved, after which a 1cm incision was made and the kidney exteriorized, the capsule incised, and the donor thymi (both lobes) transplanted in subcapsular fashion, after which the capsular defect was cauterized, and the kidney replaced intraabdominally. The flank incision was sutured closed with 6-0 Monocryl. Mice were recovered on covered warming pads with buprenorphine/flunixin analgesia, and carefully monitored during recovery.

Two weeks after the procedure, experimental NOD thymic swap mice were sublethally irradiated with 900 rads (600/300), and donor bone marrow was prepared from NOD 8.3 Thy1.1 mice by complement depletion of CD4+ and CD8+ T cells (anti-CD4 clone GK1.5, anti-CD8 clone YTS 164.5). 5e6 BM cells were transferred per recipient via tail-vein injection. Mice were analyzed by flow cytometry at eight weeks post-reconstitution or followed for diabetes incidence by weekly urine dipstick, or at more frequent intervals if they developed wasting or polyuria.

### RANK-ligand blockade

Anti-RANKL antibody (Clone IK22/5, BioXCell) or isotype control antibody (clone 2A3, BioXCell) was administered at 100μg/mouse in PBS every other day via intraperitoneal (i.p.) injections for a total of 3 injections.

### Mouse lymph node processing and purification

Mouse secondary lymphoid organs (cervical, brachial, axillary, inguinal, mesenteric lymph nodes) were procured and pooled in digestion media consisting of RPMI with 2% FBS (Sigma), with 100ug/ml DNase (Roche) and 50ug/ml Liberase (Roche), minced and agitated at 37C with gentle agitation for 30 minutes, and passed through a 70-micron filter. Cells were resuspended for magnetic column enrichment (Miltenyi LD column depletion with Streptavidin or anti-biotin microbeads and biotinylated antibodies against B220, Ter119, TCRb, +/- CD3e). Cells were then either processed directly for 10x single-cell analysis (WT LN) or sorted by flow cytometry for all live, GFP+ cells, along with WT (non-transgenic) controls for MFI thresholding. Cells were sorted into PBS with 0.04% BSA. Cell viability and counts were evaluated with ViCell XR (company name here), and samples with viability >85% used for sequencing.

### Thymus processing for flow cytometry

Mouse thymi were isolated and minced with razor blades, transferred into 15ml falcon tubes in total 10ml of DMEM (Sigma) with 2% FBS, and vortexed briefly for 15 seconds. After settling for 5 minutes, the supernatant media was removed and replaced with 4 ml of digestion media consisting of DMEM with 2% FBS (Sigma), with 100ug/ml DNase (Roche) and 50ug/ml Liberase (Roche). Thymi were digested in 37°C water bath with mechanical aid by pipetting through a glass Pasteur pipette every 4 minutes. Every 8 minutes, tubes were spun briefly to pellet undigested fragments and the digested supernatant was transferred to 20 ml of 0.5% BSA (Sigma-Aldrich), 2 mM EDTA (TekNova), in PBS (MACS buffer). This procedure was repeated for three rounds with 4ml of fresh digestion media each time until there were no remaining visible tissue fragments (total 24 minutes). The single-cell suspension was then pelleted and washed once in MACS Buffer and passed through a 70-micron filter. To enrich for stromal cells, single cell suspension of thymi were subjected to density-gradient centrifugation with a three-layer Percoll gradient (GE Healthcare) with gravities of 1.115, 1.065, and 1.0. Stromal cells are isolated from the Percoll-light fraction, between the 1.065 and 1.0 layers, and resuspended in PBS before counting and flow staining.

### scRNA-seq

ScRNA-seq libraries were generated using the 10x Chromium Controller and the Chromium Single Cell 3′ Library Construction Kit v3 according to the manufacturer’s instructions. Briefly, the suspended cells were loaded on a Chromium controller Single-Cell Instrument to generate single-cell Gel Bead-In-Emulsions (GEMs) followed by reverse transcription and sample indexing using a C1000 Touch Thermal cycler with 96-Deep Well Reaction Module (BioRad). After breaking the GEMs, the barcoded cDNA was purified and amplified, followed by fragmenting, A-tailing and ligation with adaptors. Finally, PCR amplification was performed to enable sample indexing and enrichment of scRNA-Seq libraries. The final libraries were quantified using a Qubit dsDNA HS Assay kit (Invitrogen) and a High Sensitivity DNA chip run on a Bioanalyzer 2100 system (Agilent). All libraries were sequenced using Nova-seq S4 (Illumina) with paired-end reads (2 x 150 cycles), and per-sample sequencing libraries were demultiplexed using CellRanger v5.

### ATAC with Selected Antigen profiling by sequencing (ASAP-seq)

Cells were stained with a 198 Total-seq A-conjugated antibody panel for murine (BioLegend, see Table S4 for a list of antibodies, clones and barcodes used for ASAP-seq) as previously described (57) and outlined online: https://cite-seq.com/asapseq/. Briefly, following sorting, cells were fixed in 1% formaldehyde and processed as described for the mtscATAC-seq workflow (25), with the modification that during the barcoding reaction, 0.5µl of 1µM bridge oligo A (BOA for TSA) was added to the barcoding mix. For GEM incubation the standard thermocycler conditions were used as described by 10x Genomics for scATAC-seq. Silane bead elution and SPRI cleanup steps were modified as described to generate the indexed protein tag library. The final libraries were quantified using a Qubit dsDNA HS Assay kit (Invitrogen) and a High Sensitivity DNA chip run on a Bioanalyzer 2100 system (Agilent). All libraries were sequenced using Nova-seq S4 (Illumina) with paired-end reads (2 x 150 cycles), and per-sample sequencing libraries were demultiplexed using CellRanger-ATAC v1.2.

### scRNA-seq analyses

For each sequencing library, per-cell, per-gene counts were generated from raw sequencing reads (.fastq files) using CellRanger v5 count using the mm10 reference genome. Each library (GFP and WT) yielded high quality data, including 11,697 and 10,953 cells for the GFP and WT libraries with a median number of genes detected per cell of 1,831 and 1,451, respectively. To perform integrative downstream analyses, we utilized the Seurat analytical toolkit (58). First, we filtered putative cell doublets using DoubletFinder (59) and down sampled the GFP to 3,000 cells to allow for a more balanced representation of the LN populations in the reduced dimension space. Next, we filtered for cells meeting the following criteria: greater than 500 genes detected and 1000 UMIs and no more than 10% mitochondrial RNA UMIs and no more than 15,000 total UMIs detected per cell—thresholds based on inspection of the empirical density of cells— resulting in 2,532 GFP+ cells and 6,973 WT cells. After normalization and variable gene inference, we corrected the top 30 principal components for the sequencing batch using Harmony (60). The Harmony-corrected components were then used for identifying cell neighbors, clusters, and reduced dimensionality coordinates. Visualization of populations and gene abundances were performed using the default Seurat functionality. For GFP positive analyses (Figure 3), we utilized similar thresholds but did not down sample the library to increase power to discriminate features between JCs and migDCs. RNA velocity analyses were performed using velocyto using the default parameters for this GFP+ population (41).

### ASAP-seq analyses

For this multi-modal capture, scATAC-seq data was aligned and processed using the CellRanger-ATAC v1.2 pipeline with the mm10 reference genome. For the ASAP-seq antibody tag data, per-cell, per-antibody tag counts were enumerated via the kite | kallisto | bustools framework accounting for unique bridging events as previously described (57). Cells called by the CellRanger-ATAC knee call were further filtered based on abundance accessible chromatin (>3,000 nuclear fragments) as well as accessible chromatin enrichment (>4 TSS enrichment score) and minimal non-specific antibody binding (<2000 total antibody tag counts). We further down sampled the GFP positive population to ensure a more balanced representation of all populations in the LN as described in the scRNA-seq analysis section. Next, gene activity score calculations (via Signac (61)) were performed using the label transfer functionality originally described in Seurat v3 (58) for our well-defined cluster annotations from the scRNA-seq data. We excluded cells with a <60% confidence for the maximum population assigned. Downstream analyses, including dimensionality reduction, transcription factor binding analysis, accessible chromatin visualization was all performed using functions from the Signac package (61), including those that wrap chromVAR (62).

### Cell type annotation/ImmGen similarity scores

In order to assign identities to the resulting cell subpopulations/clusters in an unbiased way, we developed an approach to query the similarity of single-cell RNA-seq profiles against all 178 publicly available reference microarray profiles in the Immunological Genome Project (ImmGen) Database (23); all ImmGen primary data were downloaded from haemosphere (https://www.haemosphere.org/datasets/show). The similarity measure we used is based on the cosine distance between appropriately normalized microarray/gene expression profiles - considering only genes (n=2,000) that were identified as highly variable in scRNA-seq. For each reference subpopulation, the corresponding ImmGen microarray profile was normalized to unit sum. The single cell gene expression profiles were first scaled to 1e4 total counts and logarithmized before being normalized to unit sum. Each expression dataset was then subsequently scaled. For each single cell profile, the resulting cosine similarities to all ImmGen profiles were further min-max scaled in a range between 0 and 1, to obtain the final ImmGen similarity scores. We applied this approach both for scoring individual cells (t-SNE visualization) and pseudo-bulk cluster profiles (clustered heatmap visualization). Final cell type annotation was carried out in a semi-supervised way, cross-validating the resulting ImmGen scores with subpopulation-specific gene expression of known markers and shown in Figure S2. An analogous workflow was devised for the ATAC scoring of populations where the top ∼9,000 variable peaks were used as a feature set before performing similarity estimates between the pseudo-bulk and single-cell ATAC-seq data generated here and the bulk ImmGen ATAC-seq database.

### Tissue-specific antigen assignment, annotation, and expression analysis

To examine tissue specific antigens (TSA), we downloaded the corresponding list (Supplemental Table 2) from Sansom et al. (32), comprising of 6,610 genes, 202 of which were significantly up-regulated in JCs compared to other Aire-expressing populations. To map these TSAs to their corresponding tissue, we downloaded the GeneAtlas MOE430 dataset from BioGPS and excluded cell lines from the data resource, resulting in 89 tissues for further examination. The distribution of tissues with JC-specific TSAs represented the tissue with the maximum expression per-gene from this resource.

### Merging and Analysis of Previously Published mTEC scRNAseq datasets

Raw transcript data from four previously published thymic scRNAseq datasets were collected and merged, filtering on genes that were detected across all four datasets (44–47) and on cells isolated from wild type mice. Data likely to correspond to doublets were excluded by filtering out cells with total reads and number of genes in the top 1 percent. Further quality control was performed by excluding cells containing fewer than 3 genes. Batch correction and normalization of raw read counts were performed with scVI (43) using the default parameters for model training and batch keys corresponding to each sequencing run within the four datasets. An scVI model trained on the top 5,000 highly variable genes was used to generate a UMAP for the merged data, which was then subset on the 18,349 cells in mTEC clusters based on cell type labels assigned in the published analysis of the individual datasets. A new UMAP was generated for this subset of the filtered data and an scVI model trained using all 16,786 genes in the dataset was used to normalize gene expression for visualization.

### Statistical Analyses and Visualization

All data points are shown in the graphs as mean value indicated and individual data points as shown. For all boxplots: center line, median; box limits, first and third quartiles; whiskers, 1.5× interquartile range. All statistical analyses were performed using GraphPad Prism 8 and R v4.0.3. Statistical significance was calculated using one-way ANOVA with Tukey multiple comparison, the unpaired two-sided t-test, or the non-parametric Mann-Whitney U test. Diabetes-free survival was analyzed using the log-rank test. *p < 0.05, **p < 0.005, *** p < 0.0005 and **** p < 0.00005 were considered significant. Experimental graphics were generated with Biorender (https://biorender.com/).

## SUPPLEMENTAL FIGURES

**S1.**
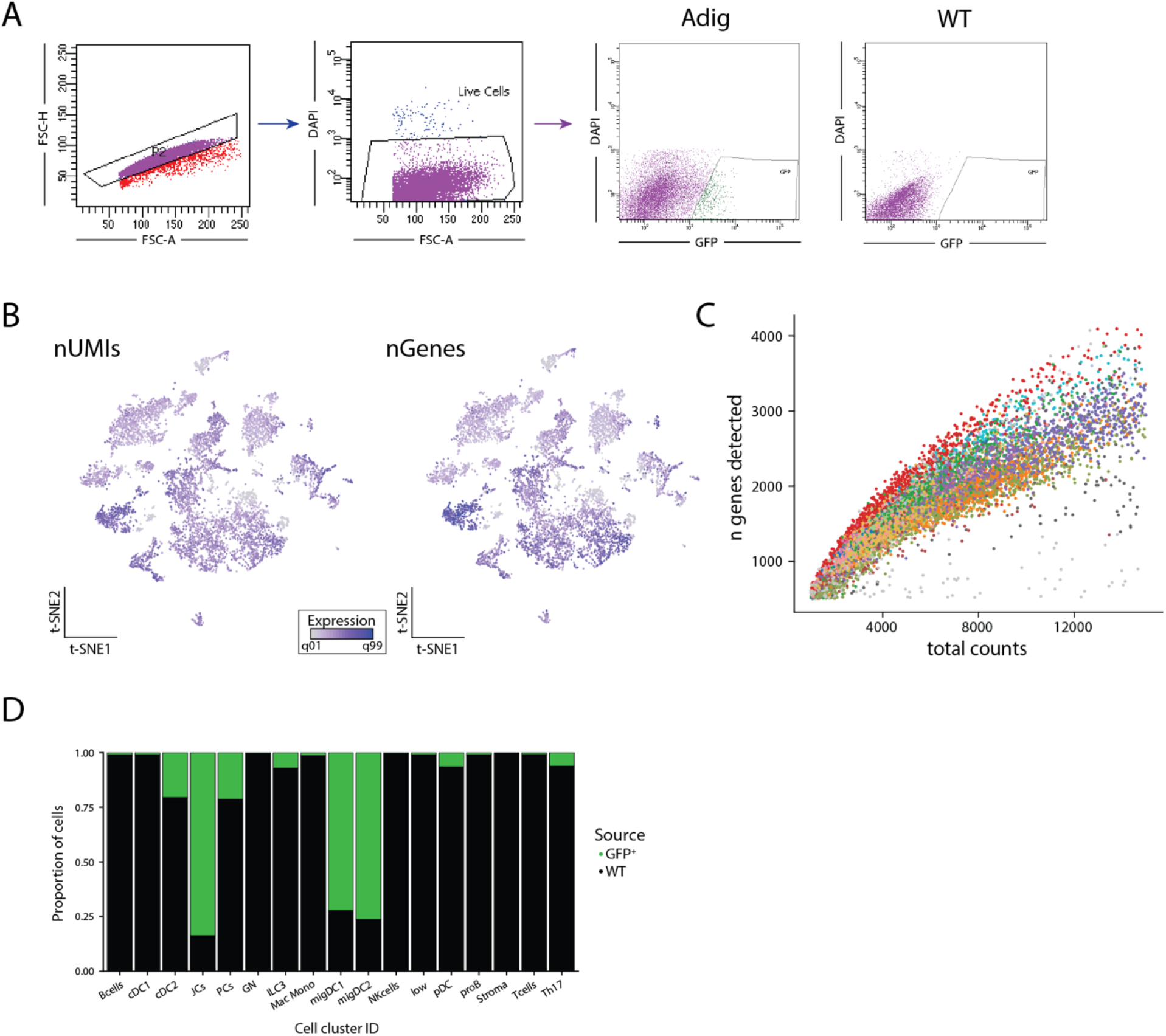
(A) Sorting strategy for single, live, GFP+ cells from Adig lymph nodes, with WT as GFP-negative control. (B) Summary of scRNA-seq quality control depicting number of unique molecular identifiers (UMIs; left) and genes (right) detected. (C) Scatter plot depicting genes and counts detected per cell. Colors represent cell clusters. (D) Proportion of cells per cluster from the GFP+ and wildtype (WT) mice lymph nodes.

**S2.**
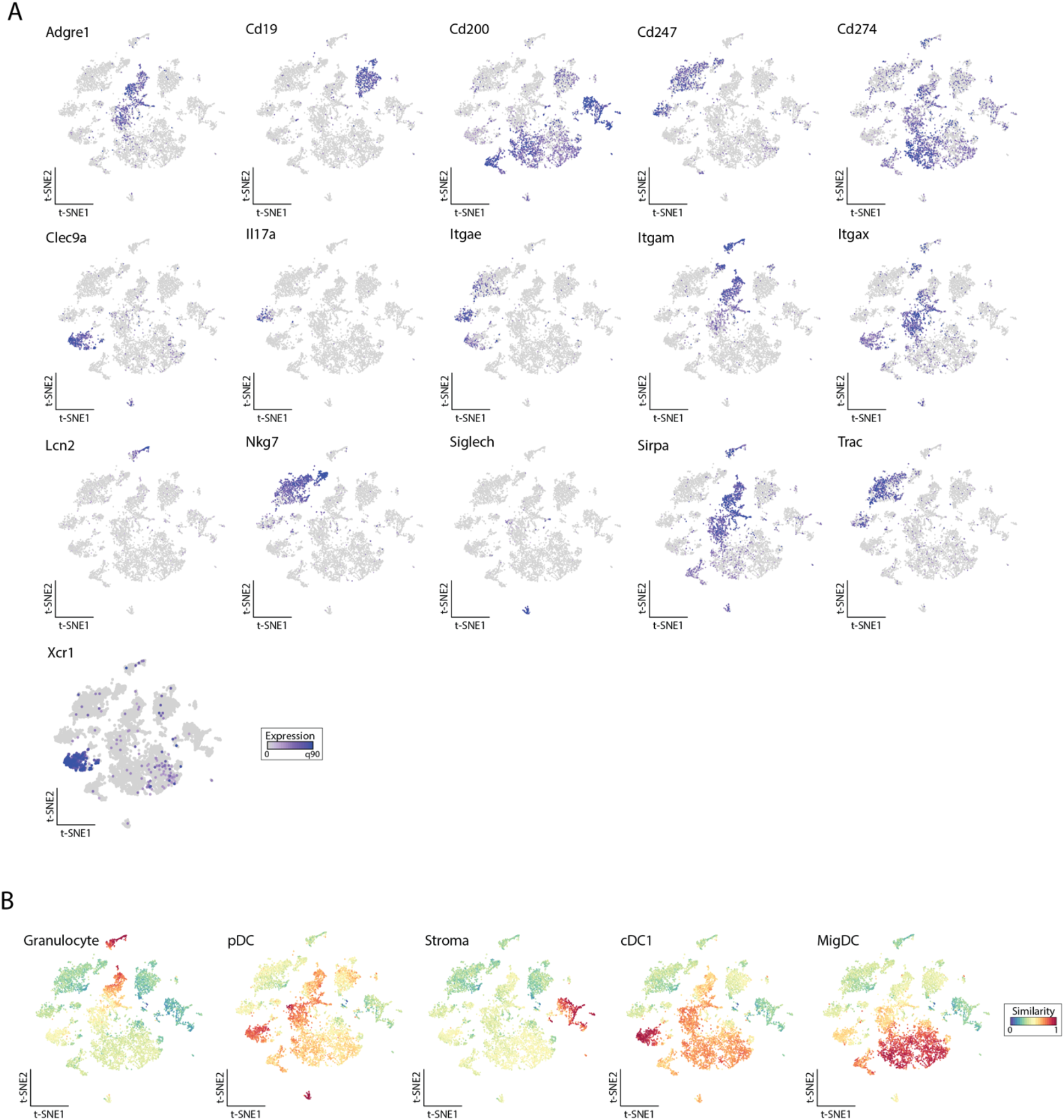
(A) Feature maps of selected individual genes expressed in merged WT/GFP+ scRNA-seq tSNE space used to assign cluster identity. (B) ImmGen similarity scores supporting annotation of cell clusters from the scRNA-seq data with reference populations as indicated. cDC1 and MigDC are replicates from other ImmGen reference populations.

**S3.**
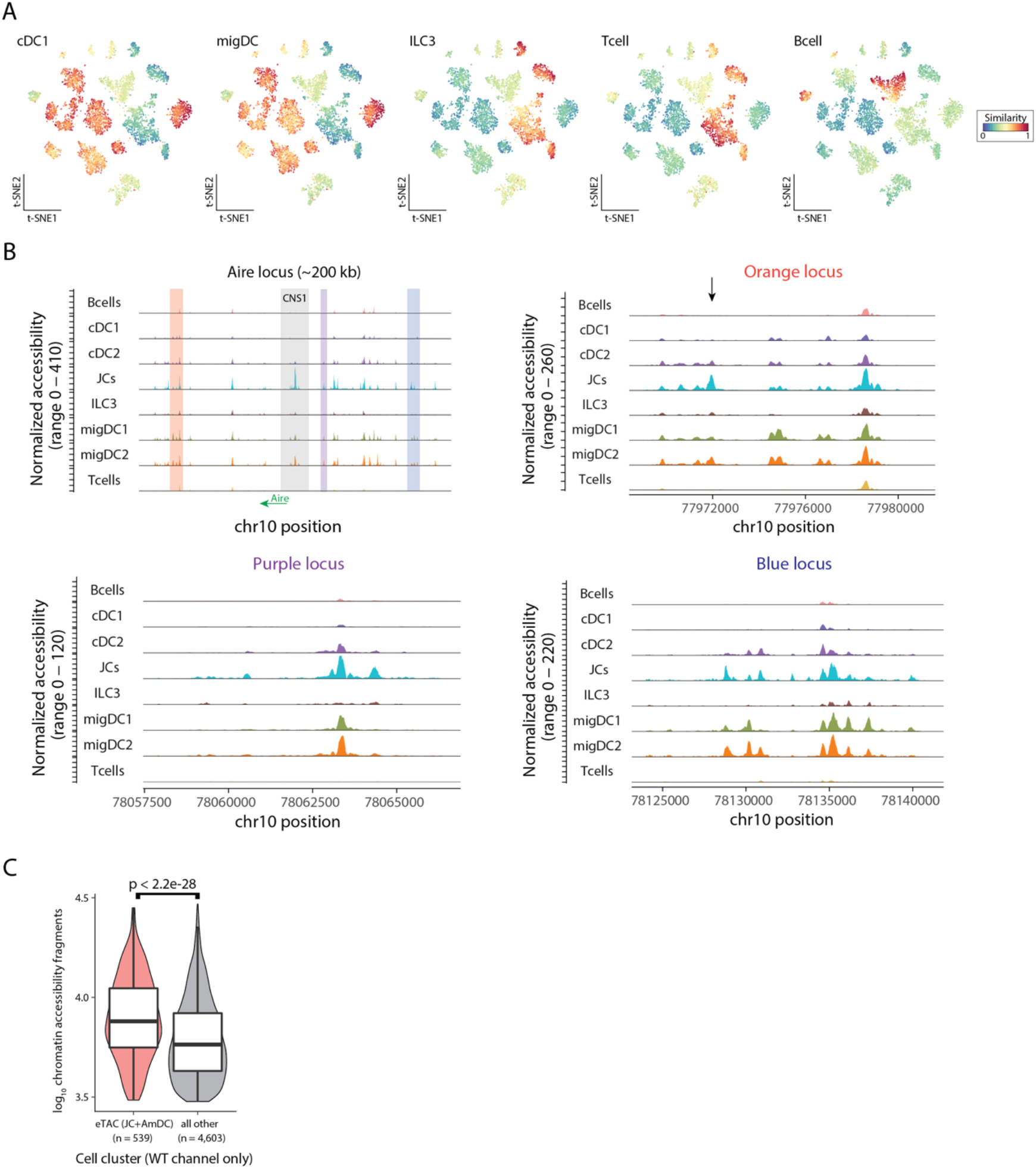
(A) ImmGen similarity scores at single-cell resolution for ATAC-seq data for selected populations. (B) Accessible chromatin landscapes near the *Aire* locus. Individual accessible regions are indicated. (C) Comparison of number of accessible chromatin fragments among only the WT sample. Boxplots: center line, median; box limits, first and third quartiles; whiskers, 1.5× interquartile range. Statistical test: Mann-Whitney test.

**S4.**
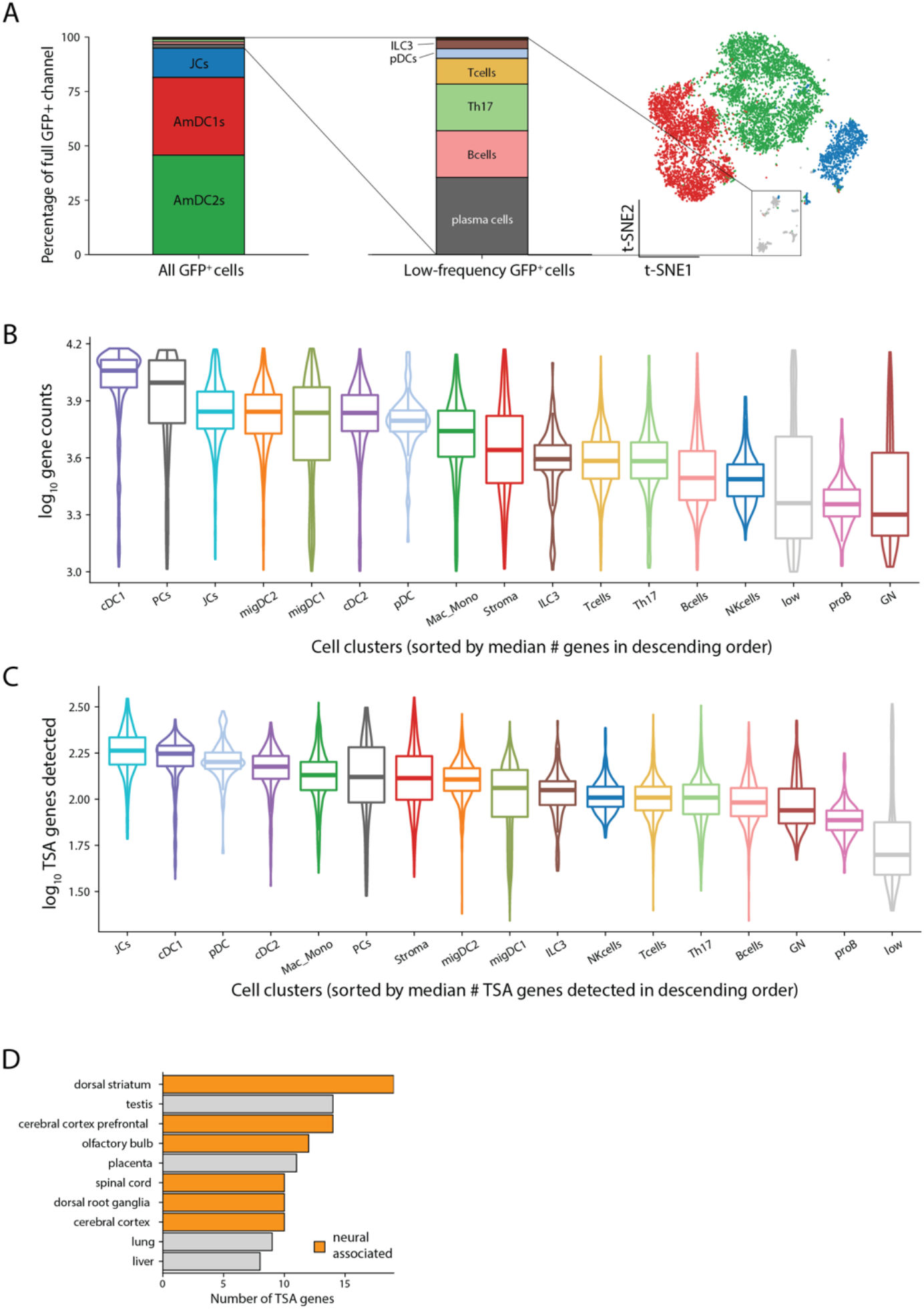
(A) Description of GFP+ populations, including low abundance populations (grey). (B) Number of genes detected and (C) number of TSAs per population. Boxplots: center line, median; box limits, first and third quartiles; whiskers, 1.5× interquartile range. (D) Count of tissue specific antigen (TSA) genes overexpressed in JCs by tissue as assigned by GeneAtlas. Neuronal tissues indicated in orange.

**S5.**
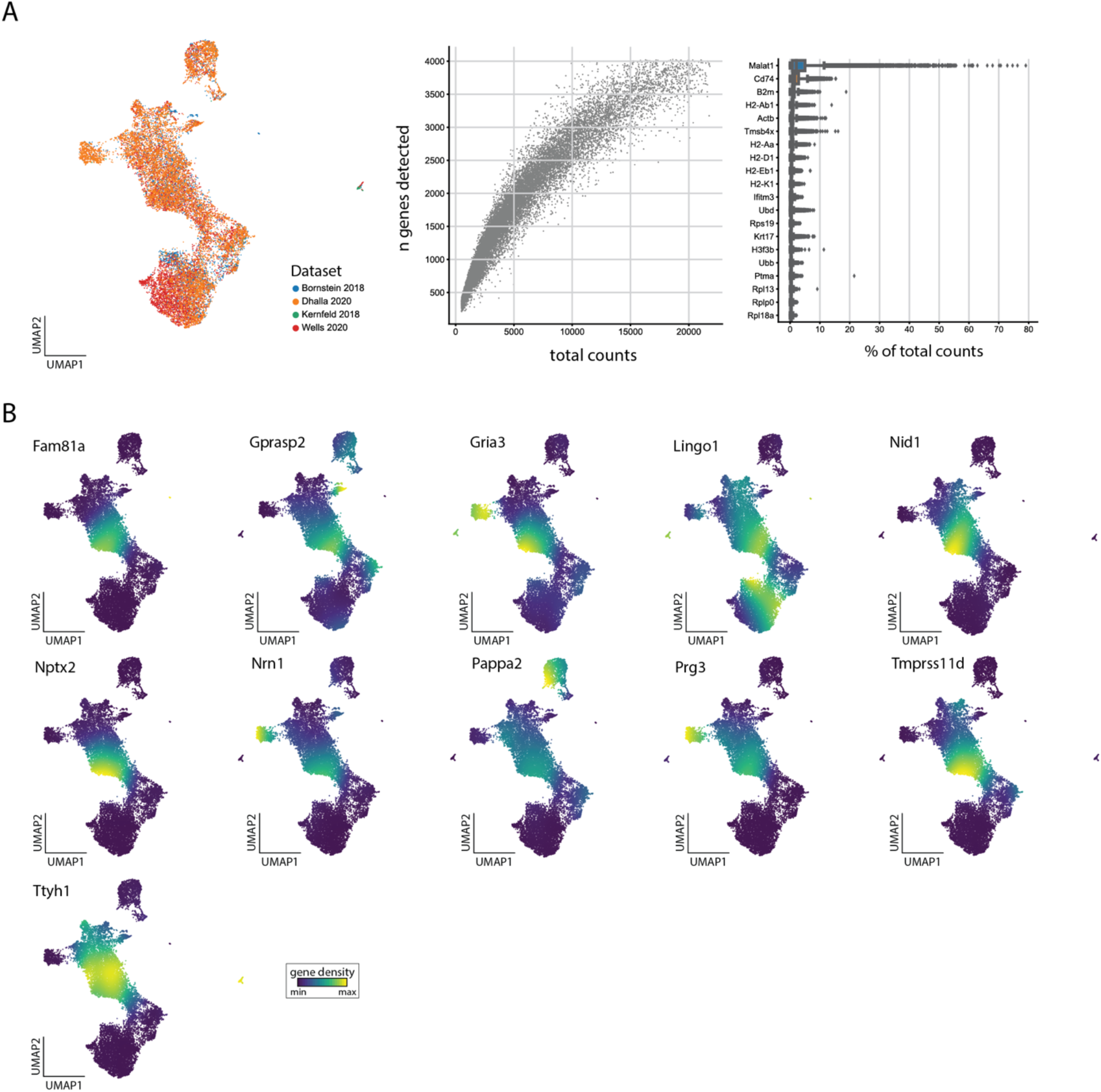
(A) Summary of quality-control for merged thymus scRNA-seq datasets. (B) Gene-expression densities of JC-specific genes in reduced dimension space from merged previously published thymic epithelial scRNAseq data.

## SUPPLEMENTAL TABLES

**S1. Summary statistics of differentially expressed genes between JCs and amDCs.**

**S2. Summary statistics of genes most over-expressed in JCs.**

**S3. Top transcription factor activity scores for JCs.**

**S4. Details of ASAP-seq surface panel.**

## Acknowledgments

We thank members of the Bluestone and Anderson labs for technical support and discussions including Alice Chan, Corey Miller, Jennifer Bridge, Jonah Phipps, and Irina Proekt. We thank Chrysothemis Brown and Sasha Rudensky for discussions. We thank the members of the UCSF Flow Cytometry Core Facility for assistance with cell sorting. **Funding:** This work was supported by the ARCS Fellowship (J.W.), a Stanford Science Fellowship (C.A.L.), NIH T32 FAVOR (J.L.B.), the NIH P01AI118688 (M.S.A.), the ASTS Fellowship in Transplantation grant (J.M.G.), the UCSF Sandler Fellows PSSP grant (J.M.G.), NIH K08230188 and U01CA260852 (A.T.S.), a Cancer Research Institute Technology Impact Award (A.T.S.), and the Juvenile Diabetes Research Foundation and Lupus Research Alliance (A.T.S.). **Author Contributions:** J.M.G. and M.S.A. conceived the study. J.W., J.L.B., C.L., A.R.G., and J.M.G. designed and performed experiments and analyzed data. C.L., J.G., J.M.G., V.N. and N.C. performed and supervised computational analysis. F.X. generated and cared for mice and performed experiments; J.M.G. J.W., C.L., and A.T.S. wrote and reviewed the manuscript. All authors read and approved the manuscript. **Competing interests:** A.T.S. is a founder of Immunai and Cartography Biosciences and receives research funding from Arsenal Biosciences and Allogene Therapeutics. **Data and materials availability:** The murine sequencing data are available through the Gene Expression Omnibus under accession GSE176282 with reviewer access token: **ufohuogmtlenjkn**.

